# Histone Deacetylases (HDACs) maintain expression of the pluripotent gene network via recruitment of RNA polymerase II to coding and non-coding loci

**DOI:** 10.1101/2023.04.06.535398

**Authors:** RDW Kelly, KR Stengel, A Chandru, LC Johnson, SW Hiebert, SM Cowley

## Abstract

Histone acetylation is a dynamic modification regulated by the opposing actions of histone acetyltransferases (HATs) and histone deacetylases (HDACs). Deacetylation of histone tails results in chromatin tightening and therefore HDACs are generally regarded as transcriptional repressors. Counterintuitively, simultaneous deletion of *Hdac1* and *Hdac2* in embryonic stem cells (ESC) reduced expression of pluripotent transcription factors, *Oct4, Sox2* and *Nanog* (OSN). By shaping global histone acetylation patterns, HDACs indirectly regulate the activity of acetyl-lysine readers, such as the transcriptional activator, BRD4. We used inhibitors of HDACs and BRD4 (LBH589 and JQ1 respectively) in combination with precision nuclear run-on and sequencing (PRO-seq) to examine their roles in defining the ESC transcriptome. Both LBH589 and JQ1 caused a marked reduction in the pluripotent network. However, while JQ1 treatment induced widespread transcriptional pausing, HDAC inhibition caused a reduction in both paused and elongating polymerase, suggesting an overall reduction in polymerase recruitment. Using enhancer RNA (eRNA) expression to measure enhancer activity we found that LBH589-sensitive eRNAs were preferentially associated with super-enhancers and OSN binding sites. These findings suggest that HDAC activity is required to maintain pluripotency by regulating the OSN enhancer network via the recruitment of RNA polymerase II.

## Introduction

Acetylation of lysine residues in histone tails is an essential post-translational modification (PTM), dynamically regulated by the opposing actions of histone acetyltransferases (HATs) and histone deacetylases (HDACs; (Verdin and Ott 2014). HDACs are generally considered to be transcriptional repressors (Nagy et al. 1997; Haberland et al. 2009; Kelly and Cowley 2013), since histone deacetylation increases inter-nucleosomal affinity and the association of N-terminal tails with negatively-charged DNA, promoting chromatin compaction (Luger and Richmond 1998; Robinson et al. 2008). In addition, loss of histone acetylation prevents the binding of bromodomain-containing proteins, such as TAFII1 (Jacobson et al. 2000) and BRD4 (Kanno et al. 2004), which act as transcriptional activators. Among the 18 HDAC enzymes present within mammalian cells, HDAC1, HDAC2, and HDAC3 (class I HDACs) have well-described roles in regulating gene expression as the catalytic core of seven multi-protein complexes: SIN3, NuRD, CoREST, MiDAC, MIER and RERE (HDAC1 and HDAC2) and NCoR (HDAC3; (Millard et al. 2017). Each complex, with its unique arrangements of components, targets HDACs to specific genomic regions through the recognition of histones and interaction with transcription factors (Millard et al. 2017).

Despite a well-established role for HDACs in transcriptional repression, loss of HDAC activity, either through genetic deletion (Zupkovitz et al. 2006; Chen et al. 2013; Dovey et al. 2013; Jamaladdin et al. 2014; Stengel et al. 2017) or HDAC inhibitors (HDACi; (Karantzali et al. 2008; Baltus et al. 2009; Kim et al. 2013; Greer et al. 2015; Gryder et al. ; Baker et al. 2023), also reduces the expression of a subset of genes. In addition, HDAC activity is required for interferon-induced transcriptional initiation of the immediate-early genes (Chang et al. 2004; Sakamoto et al. 2004). In mammalian cells, HDACs are enriched at transcriptionally active enhancers, promoters and gene bodies, suggesting an alternative and non-repressive function (Wang et al. 2009; Kidder and Palmer 2012; Whyte et al. 2012). In tumour cells, HDAC inhibition reduced the activity of super-enhancers (SE) via aberrant hyperacetylation, causing down-regulation of SE-driven transcription of genes such as *Myc* (Gryder et al. 2019a; Gryder et al. 2019b). An influential study from Métivier et al. (Metivier et al. 2003) showed that activation of the *PS2* gene required cyclical acetylation/deacetylation for transcription initiation to occur. In this model, HDACs are hypothesized to function by resetting histone-acetylation at promoters between rounds of initiation (Metivier et al. 2003). In addition to initiation, HDAC activity is also required for normal transcriptional elongation (Kim et al. 2013; Greer et al. 2015). Treatment of cells with HDACi caused increased global histone acetylation levels, resulting in mis-localisation of the bromodomain-containing protein 4 (BRD4; (Kim et al. 2013; Greer et al. 2015), an essential co-factor in the conversion of RNA polymerase II from a paused-to-elongating state (Jang et al. 2005; Yang et al. 2005; Kanno et al. 2014; Arnold et al. 2021). These observations indicate HDACs positively regulate transcription through the precise regulation of histone acetylation patterns.

Embryonic stem cells (ESCs) are widely used to study the transcriptional networks required for pluripotency and tissue-specific differentiation in early embryos (reviewed in (Martello and Smith 2014)). We have previously shown an essential role for HDAC1 and HDAC2 (HDAC1/2) in maintaining the expression of the critical pluripotent transcription factors, *Oct4, Sox2 and Nanog* (OSN; (Jamaladdin et al. 2014). Similar transcriptional changes are observed in ESCs treated with inhibitors of HDAC1-11 (Karantzali et al. 2008; Baltus et al. 2009) and bromodomain inhibitors (Di Micco et al. 2014; Liu et al. 2014; Horne et al. 2015; Gonzales-Cope et al. 2016; Finley et al. 2018). In this study, we examine the molecular mechanisms by which HDACs and the Bromodomain and Extra-Terminal motif (BET) family (BRD2-4) maintain the expression of the pluripotent network. Complete (*Hdac1/2-KO*) or partial deletion (*Hdac1-KO; Hdac2-Het*) of *Hdac1/2*, but not *Hdac3*, resulted in reduced expression of *Esrrb*, *Nanog* and *Pou5f1*. To examine the direct effects of HDAC and BRD4 inhibition on the ESC transcriptome (using LBH589 and JQ1 respectively), we performed precision nuclear run-on and sequencing (PRO-seq) analysis, which provides near nucleotide resolution of multiple nascent RNA species, including protein-coding genes and enhancer RNAs (eRNA; (Kwak et al. 2013). While JQ1 treatment induced widespread transcriptional pausing of genes, HDAC inhibition caused a reduction in the levels of both paused and elongating polymerase. We also detected a significant increase in divergent promoter upstream transcripts (PROMPTs) at genes whose expression was increased by LBH589 treatment. Finally, we used our PRO-seq data to examine eRNA expression as a measure of enhancer activity in JQ1 and LBH589-treated cells. The addition of JQ1 caused a general reduction in eRNA levels, whereas HDAC inhibition preferentially reduced eRNAs associated with SEs and OSN binding sites. Together, these findings suggest that HDAC1/2 activity is required for the maintenance of pluripotency by regulating the OSN enhancer network via the optimal recruitment of RNA pol II.

## Methods

### ESC culture and stable BRD4 expressing lines

All mouse E14 ESCs, (*Hdac1/2^L/L^*, *Hdac1^L/L^*; *Hdac2 ^L/wt^* and *Hdac3^L/^*^L^) were cultured as previously described in Jamaladdin et al., 2014. Flag-tagged BRD4 cDNA was cloned into the PB-EF1-MCS-IRES-Neo cDNA (Systems Bioscience, California, USA) and transfected into 5x10^5^ E14-ESC along with transposase using Lipofectamine2000. After 24 hr post-transfection cells were cultured at low density (5000 cells/10 cm plate) and treated with 200 μg/ml G418 (Thermo Fisher, UK) in ESC cell media for 10 days. G418-resistant single colonies were selected, expanded individually and characterized for Flag expression by western blotting.

### Protein extraction and Western Blotting

Whole-cell extracts were prepared in cell lysis buffer containing 150 mM NaCl, 50 mM Tris-HCL pH 7.5, 0.5% IEGPAL and a protease inhibitor cocktail at 4 °C for 30 min. Samples were cleared at 20,000 x g for 15 min and the supernatant was used for the examination of soluble proteins. Histones were extracted from the insoluble pellet fraction using 0.2 M sulphuric acid overnight at 4 °C. Proteins (30 μg) were resolved using SDS/PAGE and nitrocellulose membranes and were probed with the appropriate antibodies (Supplementary Table 1). The Odyssey Infrared Imaging System was used to quantify protein signals using the appropriate IR-Dye conjugated secondary antibodies (LI-COR Biosciences).

### Chromatin immunoprecipitation (ChIP)

ChIP for histone modifications was processed previously as described by (Kelly et al. 2018). ChIP for transcription factors was processed by cross-linking cells with 1% formaldehyde for 10 min before being quenched in 125 mM glycine for 5 min. Cross-linked samples were then lysed at 4 °C for 5 min in X-ChIP Buffer 1 (50 mM HEPES-KOH pH7.5, 140 mM NaCl, 1 mM EDTA, 10% glycerol, 0.5% NP-40, 0.25% Triton X-100 and a protease inhibitor cocktail), before being centrifuged at 900 x g for 5 min (4 °C) and re-suspended in X-ChIP Buffer 2 (10 mM Tris-HCl pH8.0, 200 mM NaCl, 1 mM EDTA, 0.5mM EGTA and protease inhibitors) for 5 min at 4 °C. Samples were then pelleted (900 x g for 5 min at 4 °C) and re-suspended in Sonication Buffer (10 mM Tris-HCl pH8.0, 100 mM NaCl, 1 mM EDTA, 0.5% EGTA, 0.1% Na-deoxycholate, 0.5% N-laurylsarcosine, 0.1% SDS and protease inhibitors). Samples were sonicated for 4 min (20 seconds on, 40 seconds off) using a Bioruptor Pico (Diagenode). After sonication samples were cleared with 1% triton X-100 and pelleted (16000 x g for 5 min at 4 °C). Chromatin was immunoprecipitated using 8 μg anti-H3K27ac (Catalogue No: 39133; Active Motif), 10 μL anti-BRD4 (Bethyl Laboratories, USA) or 10 μL anti-RNA pol II (Active Motif 39097) conjugated to 50 μL Dynabeads Protein G overnight at 4 °C. A list of the antibodies used can be found in Supplementary Table 1. Beads were washed for 5 min in wash buffer A (50 mM Tris-HCl, pH 7.5, 10 mM EDTA, 75 mM NaCl), wash buffer B (50 mM Tris-HCl, pH 7.5, 10 mM EDTA, 125 mM NaCl), and wash buffer C (50 mM Tris-HCl, pH 7.5, 10 mM EDTA, 175 mM NaCl). DNA was purified using IPure kit v2 (Diagenode, Belgium) according to the manufacturer’s instructions.

### RNA extraction and quantitative Polymerase chain reaction (qPCR)

Total RNA was isolated using TRIzol (Thermo Fisher, UK) and Direct-zol™ RNA Kit (Zymo Research, UK) according to the manufacturer’s conditions. Samples were treated with RNase to remove genomic contamination. 500 ng of RNA was reverse-transcribed using 4 μL Q-Script cDNA SuperMix (Quanta Bioscience) in a 20 μL reaction according to the manufacturer’s conditions. cDNA was diluted 1 in 5 with DNase and RNase-free water. For each qPCR reaction, 2 μL of diluted cDNA or ChIP-DNA, was amplified in 10 μL reactions containing 2x SensiMix (Bioline) and 1 μM of forward and reverse primers. A complete list of primer pairs, sequences and annealing temperatures used in the study can be found in Supplementary Table 2. Reactions were carried out on a BioRad CFX-Connect under the following conditions: initial denaturation at 94 °C for 10 min, followed by 40 cycles of 94 °C for 10 s, primer-specific annealing for 20 s, and 72 °C for 5 s. Primer efficiency was calculated using absolute quantification from a standard curve. Where applicable statistical differences were identified using a Paired two-tailed Student t-test (Prism v7, GraphPad).

### Microarray analysis

Comparative gene expression profiles were generated using the Illumina mouseWG-6, v2 (LBH589 and *Hdac1^L/L^; Hdac2^L/wt^*) or Agilent SurePrint G3 Mouse Gene Expression v2 8x60K microarray (*Hdac3^L/L^*) according to manufacturer’s instructions. Quality control of total mRNA was performed using a 2100 Bioanalyser (Agilent) and only RNA integrity numbers (RIN) of 8 or higher were selected for processing and array hybridization. LBH589 and *Hdac1^L/L^; Hdac2^L/wt^* arrays were analysed using Partek Genomics Suite (Partek, UK). Gene set enrichment analysis (GSEA; (Subramanian et al. 2005) was performed using GSEA software, version 2. Data files from the analysis have been deposited in the Gene Expression Omnibus (GEO) database (GSE136620 and GSE136618 as part of the SuperSeries GSE136621).

### Precision nuclear run-on assay (Pro-seq)

Nuclear run-on and library construction were performed as previously described (Kwak et al. 2013; Zhao et al. 2016). Briefly, 2x10^7^ nuclei were isolated following drug treatments carried out in biological replicates. Nuclear run-ons were carried out in the presence of 375 μM GTP, ATP, UTP and biotin-11-CTP with 0.5% Sarkosyl at 30 °C for 3 min. Total RNA was isolated by Trizol extraction, hydrolyzed in 0.2N NaOH, and nascent RNAs were isolated by Streptavidin bead binding (Dynabeads® MyOneTM Streptavidin T1 magnetic beads, ThermoFisher Scientific). Following 3’ adapter ligation, 5’ cap removal and triphosphate repair, 5’ hydroxyl repair, and 5’ adapter ligation, RNA was reverse transcribed and libraries were amplified. Following PAGE purification, PRO-seq libraries were provided to Vanderbilt Technologies for Advanced Genomics (VANTAGE) for sequencing (Illumina Nextseq500, SE-75).

Analysis of PRO-seq data, including eRNA analysis, was performed using the Nascent RNA Sequencing Analysis (NRSA) pipeline employing alignment files in the bam format as input (Wang et al. 2018). Adapter trimming was conducted with Trimommatic-0.32 (Bolger et al. 2014) before alignment to mm10 (*mus musculus)* genome build using Bowtie 2 (Langmead and Salzberg 2012). All reads less than 10 bp and ribosomal RNA reads were moved prior to inputting bam files in to the NRSA pipeline. Transcriptional rates for genes were determined as follows: promoter-proximal regions were determining by identifying the largest number of reads in a 50 bp window within ±500 bp from known TSSs; gene body regions were identified within + 1 kb downstream of a TSS its transcription termination site (TTS); and pausing index were calculated by the ratio of promoter-proximal read density over gene-body read density. For enhancer analysis, NRSA identifies intergenic enhancers as paired bi-directional transcripts having a gap shorter than 400 bp between their 5’ ends. Enhancer centres are defined as the midpoints of 5’ end pairs. Transcripts within +/-2Kb of any annotated RefSeq gene are excluded from enhancer calling. Downregulated eRNA overlapping with mouse ESC super-enhancers regions were identify using the overlapBed function in Bedtools. Bed files and coordinates for mouse ESC super-enhancers were converted to mm10 genome build using the UCSC lift over tool.

Metagene plots of nascent RNA levels around regions of interest were generated using the annotatePeaks.pl function in HOMER (http://homer.ucsd.edu/homer/index.html,v4.9,2-20-2017). OCT4-SOX2-NANOG (OSN) binding sites were downloaded from and converted to mm10 genome build using the UCSC lift over tool. Coordinates for transcription start site (TSS) of pluripotent genes or, genes up-regulated following LBH589 treatment, were obtained from UCSC genome (https://genome.ucsc.edu). Average Pro-Seq reads around these regions of interest (TSS±2kb or 5kb) were determined at 25 bp resolution and plotted as histograms in GraphPad Prism version 7 (https://www.graphpad.com/scientific-software/prism/). Data files from the analysis have been deposited at the Gene Expression Omnibus (GEO) database (GSE136860 as part of the SuperSeries GSE136621).

### Super-enhancer analysis

Mouse ESC super-enhancers (Whyte et al. 2013) and OSN binding sites (Chen et al. 2008) were converted to the mm10 genome build using the UCSC lift-over tool. Bedtools were then used to identify overlap between super-enhancers and changes in eRNA levels following either LBH589 or JQ1 treatments.

## Results

### HDAC1 and 2 (HDAC1/2) maintain pluripotency in a gene dosage-dependent manner

We previously showed that conditional deletion of either *Hdac1* or *Hdac2* alone does not alter the expression of pluripotent transcription factors in the mouse embryonic stem cells (ESCs; (Dovey et al. 2010b). In contrast, double knockout (DKO) of *Hdac1* and *Hdac2* caused a reduction in *Pou5f1* and *Nanog*, before a catastrophic loss of cell viability (Jamaladdin et al. 2014). ESCs that retain a single copy of *Hdac2* (*Hdac1-KO; Hdac2-Het*) on the other hand, are viable but have a significant reduction in the total deacetylase activity (Jamaladdin et al. 2014). To analyse the transcriptome in ESCs with differing levels of HDAC1/2 activity, we performed a comparative microarray analysis of DKO and *Hdac1-KO; Hdac2-Het* cells at 0 (Ctrl) and 3 days post deletion. While there were numerous upregulated transcripts following KO, consistent with a role in gene repression, there were also many genes whose activity was down-regulated by a reduction in HDAC activity. Both DKO and *Hdac1-KO; Hdac2-Het* ESCs revealed significantly (FDR<0.05) reduced expression of the pluripotency genes, *Pou5f1, Nanog, Esrrb* and *Phc1* (Figure 1A). DKO cells also showed a reduction in additional pluripotent factors, including *Klf4*, *Gdf3*, and *Sall1* (Figure 1A – compare upper and lower panels). Gene set enrichment analysis (GSEA; (Mootha et al. 2003; Subramanian et al. 2005) revealed both DKO and *Hdac1-KO; Hdac2-Het* cells exhibited a reduction in pluripotency-associated gene signatures when compared to published pluripotent networks (Figure 1B, Supplementary Figure 1B). Importantly, reduced expression of pluripotent factors was not accompanied by an increase in differentiation markers (*Gata4*, *Fgf5* or *Sox17*, Supplementary Figure 1A), indicating a direct requirement for deacetylase activity rather than a generalized response to ESC differentiation. To confirm that changes in mRNA were also reflected in protein levels we examined NANOG and POU5F1 by western blotting. Upon simultaneous deletion of *Hdac1/2* we observed a 5.5- and 3.4-fold reduction in NANOG and POU5F1 protein levels respectively (Figure 1C). In comparison, compound *Hdac1-KO; Hdac2-Het* ESCs expressed 2-fold fold less NANOG and POU5F1 (Figure 1C), a smaller reduction than DKO cells, demonstrating a gene-dosage effect of HDAC1/2 activity on pluripotent gene expression. We next asked if other Zn^2+^-dependent HDACs (HDAC1-11) influenced the expression of the pluripotent network. In mouse ESCs only HDAC1-6 are expressed at significant levels (Zhuang et al. 2018). However, HDAC4 and HDAC5 have little or no deacetylase activity (Lahm et al. 2007) and HDAC6 is cytoplasmic (Boyault et al. 2007; Chen et al. 2013), leaving only HDAC3. We, therefore, analysed pluripotent gene expression in our conditional *Hdac3* KO ESCs (Simandi et al. 2016) 3 days after deletion compared to undeleted control cells. In contrast to HDAC1/2 DKO cells, cells lacking HDAC3 showed no significant change in *Esrrb*, *Klf4*, *Nanog*, *or Pou5f1* (Supplementary Figure 1C). Taken together, these data demonstrate a positive role for HDAC1/2 in the expression of pluripotent factors in ESCs.

**Figure 1:**
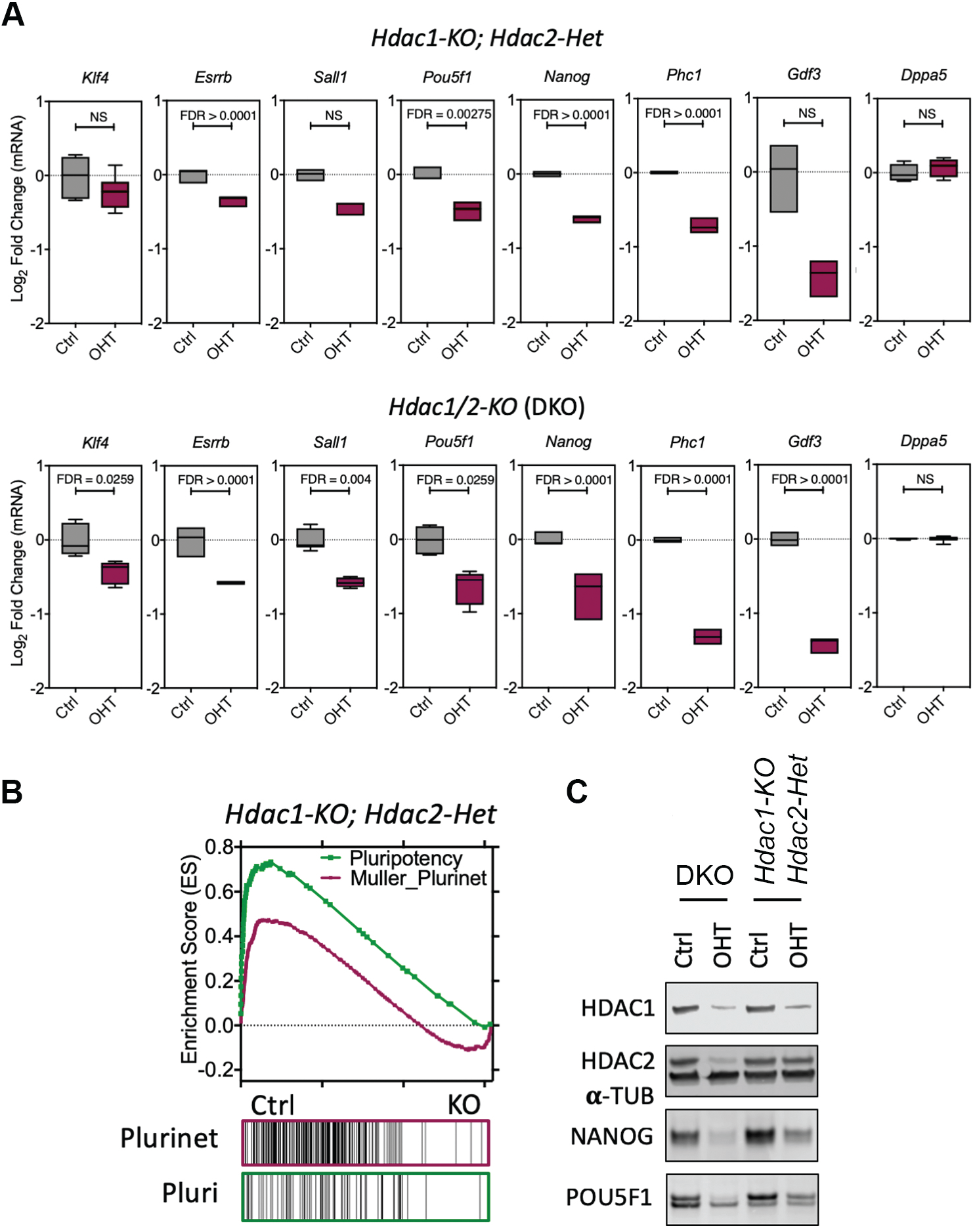
HDAC1 and 2 (HDAC1/2) maintain pluripotency in a gene dosage-dependent manner. (A) Microarray data from n = 3 biological replicates showing relative expression (log_2_ fold-change) of indicated pluripotent genes in *Hdac1/2-KO* (DKO) and *Hdac1-KO; Hdac2-Het* cells at 0 (Ctrl) and 3 days post deletion (OHT). (B) Gene set enrichment analysis (GSEA) plot of microarray data showing Ctrl samples are enriched for the Muller Plurinet (ES=0.47; P<0.0001) and pluripotency (ES=0.70; P<0.0001) gene sets compared to *Hdac1-KO; Hdac2-Het* KO cells. (C) Western blot data for HDAC1, HDAC2, NANOG and POU5F1 proteins in *Hdac1/2-KO* (DKO) and *Hdac1-KO; Hdac2-Het* cells at 0 (Ctrl) and 3 days post deletion (OHT). α-Tubulin (α-TUB) was used as a protein loading control.

To examine the role of HDAC1/2 in defining the pattern of global histone acetylation in relation to gene expression, we performed ChIP-seq and compared the levels of acetylation at histone H3 Lys27 (H3K27ac) in DKO versus control cells. H3K27ac is a marker of both active promoters and enhancers (Rada-Iglesias et al. 2011; Shen et al. 2012). We detected a robust increase in both the H3K27ac signal around transcriptional start sites in the absence of HDAC1/2 (Figure 2A, 2B), which increased yet further when we focused on genes whose expression was up-regulated in DKO cells (Figure 2C). This can be seen clearly in individual genes such as *Amn* and *Sfn,* whose mRNA levels were increased 6.8-fold and 2.9-fold, respectively (Figure 2E). At genes whose transcription is down-regulated in DKO cells, we observed relatively little change in acetylation around the promoter (Figure 2C). The pluripotent transcriptome is maintained in part through a limited number of clustered enhancers, referred to as super-enhancers (SEs), that display elevated levels of H3K27ac and transcription factor binding (e.g. MED1 and BRD4) compared to regular enhancers (Whyte et al. 2013). Surprisingly, we observed a significant increase in H3K27ac at the *Nanog* (-40 kb) SE, despite a reduction in *Nanog* mRNA levels in the absence of HDAC1/2. To examine if this was a general effect, we measured H3K27ac levels at all genomic loci designated as SEs in ESCs (Whyte et al. 2013) and found a significant increase in signal across the length of the enhancer, accentuated at the periphery (Figure 2D). These data suggest, regarding SE function, that the increased acetylation occurring in the absence of HDAC1/2 does not correlate with increased transcriptional activity, since acetylation goes up, while expression goes down (Figure 2D).

**Figure 2:**
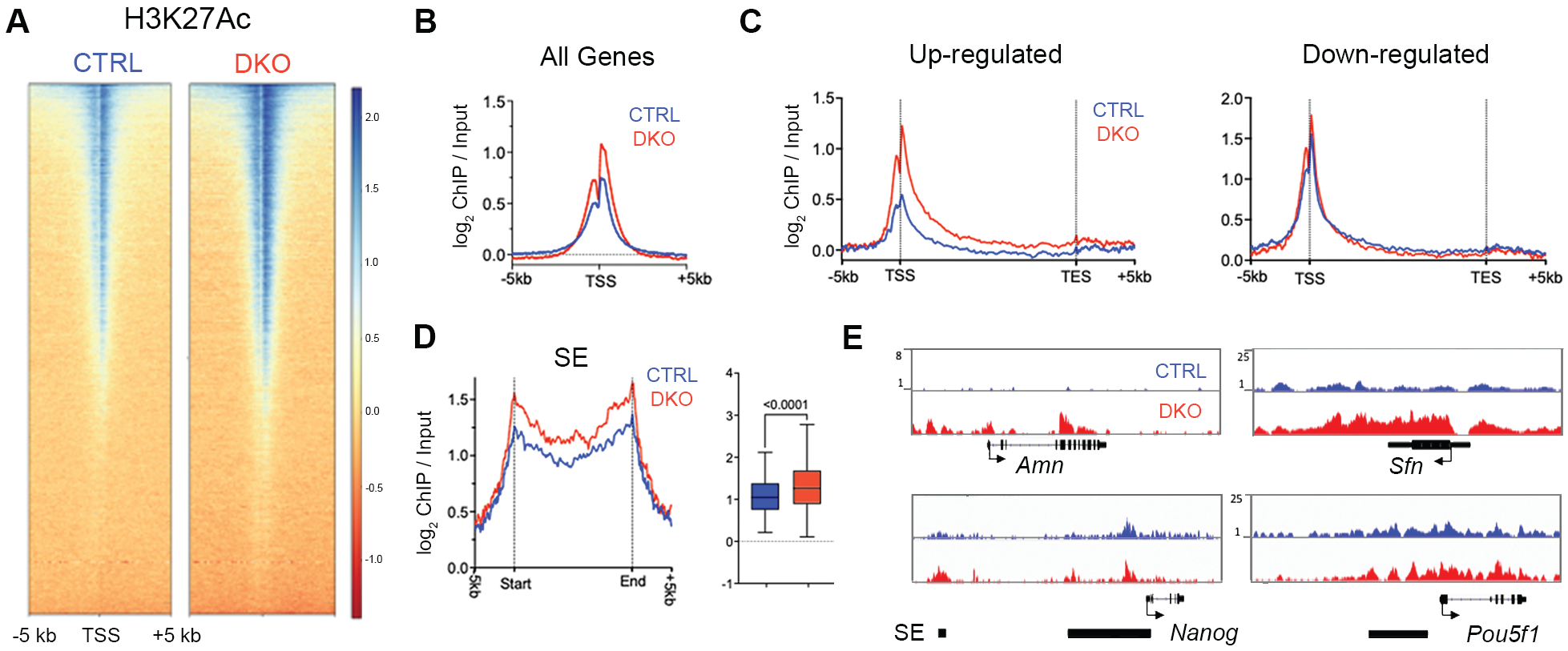
Super-enhancers have Increased histone H3 Lys27 acetylation (H3K27Ac) in the absence of HDAC1/2, but does not correlate with transcriptional activity. (A) Heatmaps show ChIP-seq data for the enrichment of H3K27ac levels (Log_2_-fold) +/- 5kb of the TSS of Refseq genes in control and *Hdac1/2* double knockout (DKO) ESCs. (B) A meta-analysis of relative H3K27ac levels relative to input for (B) all genes, (C) differentially expressed genes and (D) super-enhancer (SE) regions in control versus DKO ESCs. (E) H3K27ac levels in control versus DKO ESCs for the indicated genes.

### Acute HDAC inhibition attenuates pluripotent gene expression without inducing differentiation

Investigating the mechanistic relationship between HDAC1/2 and pluripotency using KO models is challenging because the half-lives of HDAC1/2 are approximately 24 hrs, and it takes 3-4 days until the proteins turnover completely (Jamaladdin et al. 2014). Therefore, to evaluate the direct effects of HDAC activity on gene expression in ESCs, we treated E14 cells with the HDAC inhibitor, LBH589 (panobinostat), which has been approved by the FDA and EMA to treat patients with multiple myeloma (Laubach et al. 2015). To identify HDAC-dependent immediate early genes we performed a comparative microarray analysis on mRNA isolated from ESCs treated with LBH589 at 2, 6 and 18 hours compared to untreated controls. A similar proportion of genes showed increased and decreased expression at all time points, indicating HDAC activity has both positive and negative effects on transcription in ESCs (Figure 3A). Consistent with the *Hdac1/2 DKO* model, GSEA demonstrated a reduction in pluripotency-associated gene signatures following treatment with LBH589 (Figure 3B). Importantly, germ layer differentiation was not stimulated (Figure 3C), again implying that loss of pluripotency was caused by a reduction in HDAC activity rather than as a consequence of differentiation. We observed reduced expression of *Nanog*, *Klf4* and *Sox2* as early as 2 hrs, with progressively lower expression levels at 6 and 18 hrs, respectively (Figure 3D). To confirm that these changes were a direct result of impaired transcription, and remove any confounding effects of mRNA half-life, we pulse-labelled ESCs with 5-Ethynyl Uridine (EU) to label and purify nascent mRNA, followed by RT-qPCR. In the presence of LBH589, nascent *Pou5f1* and *Nanog* transcripts decreased within 1 hr of treatment, while the expression of control genes, *Tfcp2l1* and *Thy1* was unaltered (Figure 3E). Together, these findings demonstrate that acute inhibition of HDAC activity in ESCs directly attenuates pluripotent gene expression without inducing cellular differentiation. Histone acetylation correlates with active transcription in all eukaryotes (Barnes et al. 2019) and is dynamically regulated by HDAC enzymes. We, therefore, addressed the level of H3 Lys27 acetylation (H3K27ac) at the *Nanog* and *Pou5f1* promoters and regulatory regions following HDAC inhibition using ChIP-qPCR. The addition of LBH589 stimulated a rapid increase of H3K27ac at cis-regulatory loci, including enhancers and promoters of both *Nanog* and *Pou5f1* (Figure 3F), but not at developmentally regulated genes (*Gata2* and *Spink2*), in a similar time-frame to the reduction in transcription. This is an intriguing result as it demonstrates increased histone acetylation at transcriptionally down-regulated genes, suggesting that active turnover, as part of a transcriptional acetylation/deacetylation cycle (Metivier et al. 2003), or the global pattern of histone acetylation (Greer et al. 2015), is more important than the simple accumulation of acetylation at a particular locus.

**Figure 3:**
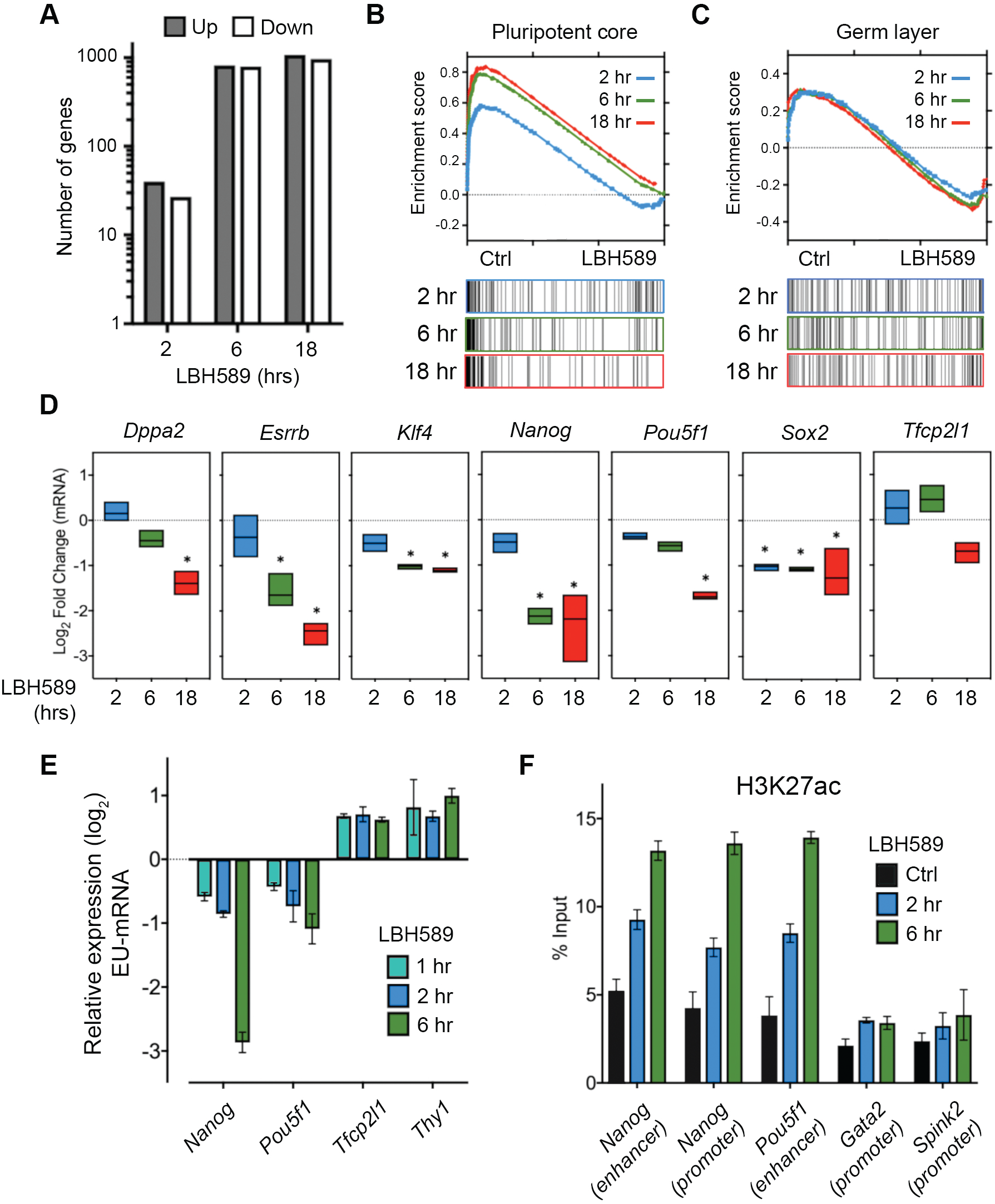
HDAC inhibition attenuates pluripotent gene expression without inducing differentiation. (A) Microarray analysis: the number of genes differentially expressed (1.5-fold; FDR ≤0.05) at the indicated times following LBH589 treatment. (B) Gene set enrichment analysis (GSEA) plot showing enrichment for the defined pluripotency gene set (Kim_Core_Module) in Control versus LBH589 treated ESCs. LBH589 2hr (ES=0.527; P<0.0001), 6hr (ES=0.789; P<0.0001) and 18hr (ES=0.83; P<0.0001). (C) GSEA for Ctrl and LBH589 samples showed equal enrichment for the formation of the primary germ layer (GO:0001704; Germ Layer). (D) Microarray data from n = 3 biological replicates showing relative expression of indicated pluripotent genes in ESCs treated with LBH589 for 2, 6 and 18hrs. The statistical difference was calculated using Benjamini & Hochberg’s false discovery rate (FDR). Asterisk (*) denotes significant changes in expression (≤1.5-fold; FDR≤0.05) relative to untreated control ESCs. (E) RT-qPCR was used to quantify nascent RNA transcription at *Nanog, Pou5f1* and *Tfcp2l1*. Values represent relative Log_2_ mean (± SEM) of n = 3 technical replicates. (F) ChIP-qPCR analysis of H3K27ac enrichment at the indicated regions. Bars represent the mean (± SEM) of n = 3 technical replicates from one immunoprecipitation.

### LBH589 and JQ1 target the same subset of pluripotent genes

It has been proposed that increasing global histone acetylation levels using HDAC inhibitors (HDACi) can sequester BRD4 away from critical sites, thereby reducing the transcriptional elongation (Greer et al. 2015). BRD4 knockdown (Liu et al. 2014) or inhibition using the bromodomain inhibitor, JQ1 (Liu et al. 2014; Horne et al. 2015; Finley et al. 2018), has been reported to reduce the expression of pluripotent factors. To explore the relationship between HDAC activity and BET family members (BRD2-4) in pluripotent gene expression, we treated ESCs with LBH589 (50 nM) and JQ1 (1 μM) for 6 and 18 hours. JQ1 treatment reduced the expression of *Nanog*, *Pou5f1*, Sox2, *Dppa3*, *Gdf3* and *Esrrb* to a similar degree and in the same time frame as LBH589, suggesting that there may be a common mode of action (Figure 4A). Therefore, to directly assess the effects of LBH589 on BRD4 recruitment we assayed the presence of BRD4 at pluripotent enhancers and promoters using ChIP-qPCR. BRD4 levels at the proximal enhancers of *Nanog* and *Pou5f1*, and the *Nanog* promoter, were reduced significantly after both JQ1 and LBH589 treatment (Figure 4B). The loss of BRD4 binding indicated an HDACi-dependent repositioning of BRD4 away from critical regulatory regions. We reasoned that if BRD4 levels were limiting *Nanog* expression, over-expression of BRD4 would rescue the effects of HDAC inhibition. To test this hypothesis, we generated ESCs stably overexpressing BRD4, and consistent with previous reports (Wu et al. 2015), observed a 4.6-fold increase in *Nanog* expression (Figure 4C). However, treatment with LBH589 caused a 2.4-fold (P<0.0001) and 1.7-fold (P=0.0241) reduction in *Nanog* levels with or without additional BRD4 respectively (Figure 4C). Taken together, these results indicate that sequestration of BRD4 can indeed limit *Nanog* expression; while the inability of additional BRD4 to rescue the effects of HDACi treatment suggests additional BRD4-independent roles for HDACs at pluripotent genes.

**Figure 4:**
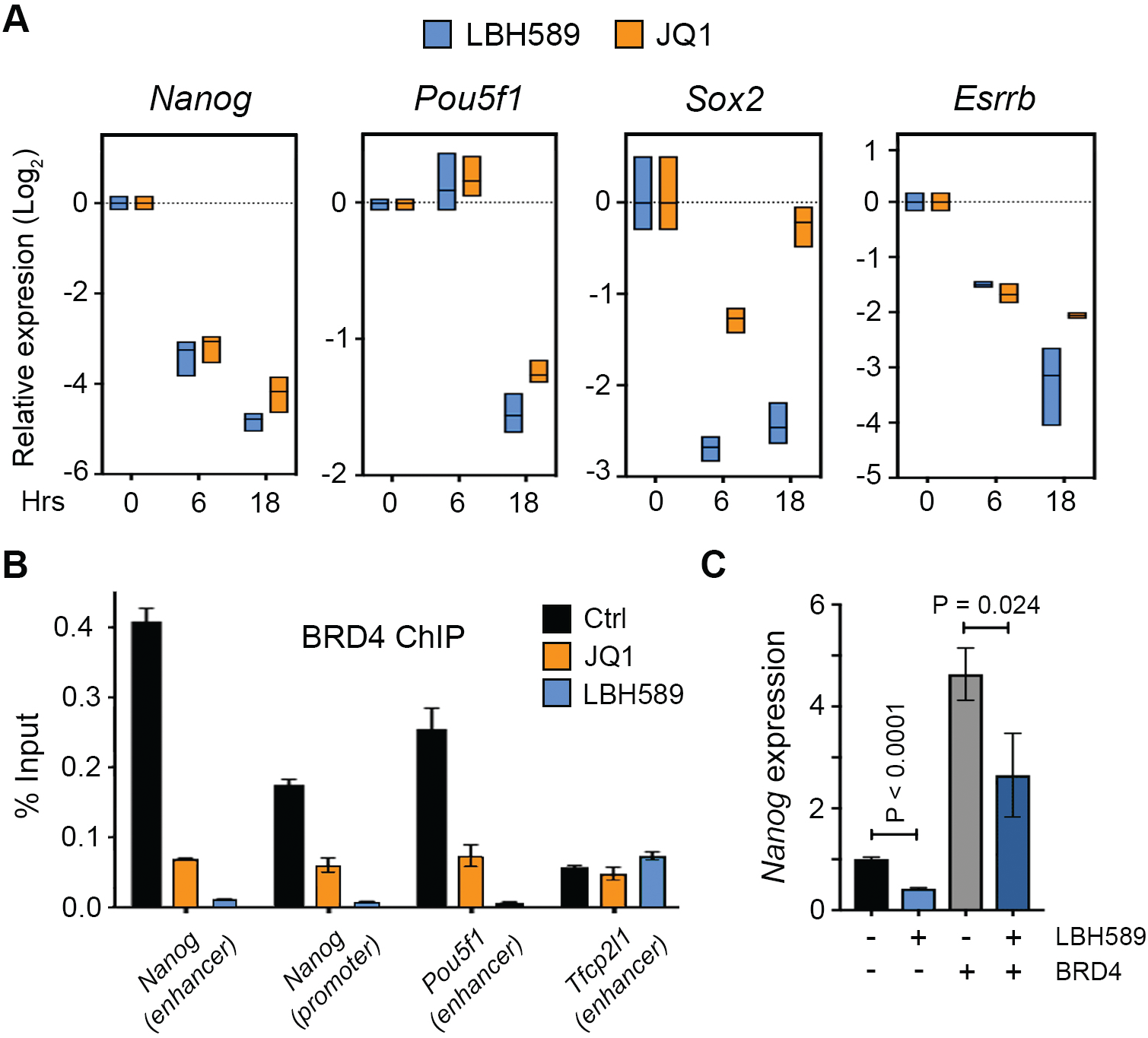
LBH589 and JQ1 target the same subset of pluripotent genes and reduce BRD4 recruitment. (A) RT-qPCR analysis of pluripotent gene expression following the treatment of ESCs with LBH589 or JQ1 for the indicated time points. Box plots represent the max-min expression range and mean from three biological replicates. (B) ChIP-qPCR analysis of BRD4 enrichment at the indicated regions. Bars represent the mean (± SEM) of n = 3 technical replicates from one immunoprecipitation. (C) RT-qPCR analysis for *Nanog* expression in ESCs treated with LBH589 and/or over-expressing BRD4, as indicated. Values represent mean (± SEM) relative expression compared to wt of n=6 biological replicates. Statistical differences were calculated using a paired Student t-test.

### HDAC inhibition caused increased expression of promoter upstream transcripts (PROMPTs) and a loss of promotor proximal polymerase

To examine the roles of HDACs and BET family members in shaping the ESC transcriptome we performed precision nuclear run-on and sequencing (PRO-seq) analysis (Kwak et al. 2013). PRO-seq enables the quantification of nascent RNAs and transcriptionally-engaged RNA polymerases across the genome; effectively an assay which combines the attributes of nascent RNA-seq with ChIP-seq for RNA polymerase II (RNA pol II). To remain consistent with previous experiments we compared PRO-seq outputs in LBH589 treated cells at 2 hr and 6 hr, and JQ1 at 2 hr. Although our study was primarily focused on pluripotent genes, we initially examined the role of HDACs from a global perspective. Analysis of RNA pol II occupancies across the transcription start sites (TSS ±5kb) of RefSeq annotated genes demonstrated significant differences in JQ1 and LBH589 treated cells (Figure 5A). BRD4 promoted the conversion of proximally-paused polymerases to an elongating form via the recruitment of P-TEFb (Jang et al. 2005; Yang et al. 2005). We observed a similar mechanism in our system, as 64% of JQ1-sensitive genes showed an increase in pausing (clusters A, B and C) (Figure 5B). Since LBH589 and JQ1 reduced the expression of pluripotent genes (Figure 4A), our expectation was that loss of deacetylase activity would affect RNA Pol II pausing in a similar fashion to JQ1. However, in contrast, the majority of genes in LBH589 treated cells showed a reduction in paused polymerase (clusters D, E and F) (Figure 5B). These data imply fundamental mechanistic differences in HDAC and BET family function. We also observed increased transcript levels upstream of the TSS in LBH589 treated cells, particularly at genes displaying increased gene body transcription (Figure 5A and 5B; clusters C and D). In mammalian cells, active promoters generate short and unstable promoter-upstream transcripts (PROMPTs) on the opposite strand to protein-coding genes (Preker et al. 2008). To examine the effects of HDAC inhibition on the generation of PROMPTS we focussed on genes whose expression is up-regulated by LBH589 treatment (clusters C and D). Metagene plots demonstrate that LBH589, but not JQ1, produced an increase in PROMPT levels (Figure 5C). Examination of individual up-regulated genes, *Fos*, *Tpst2* (Figure 5D), *Plat* and *Nrip3,* emphasises the level of increased transcription of PROMPTs following HDAC inhibition. These findings imply HDACs play a role in reducing PROMPT generation, either directly through RNA pol II recruitment, or indirectly via histone acetylation levels in the vicinity of the TSS.

**Figure 5:**
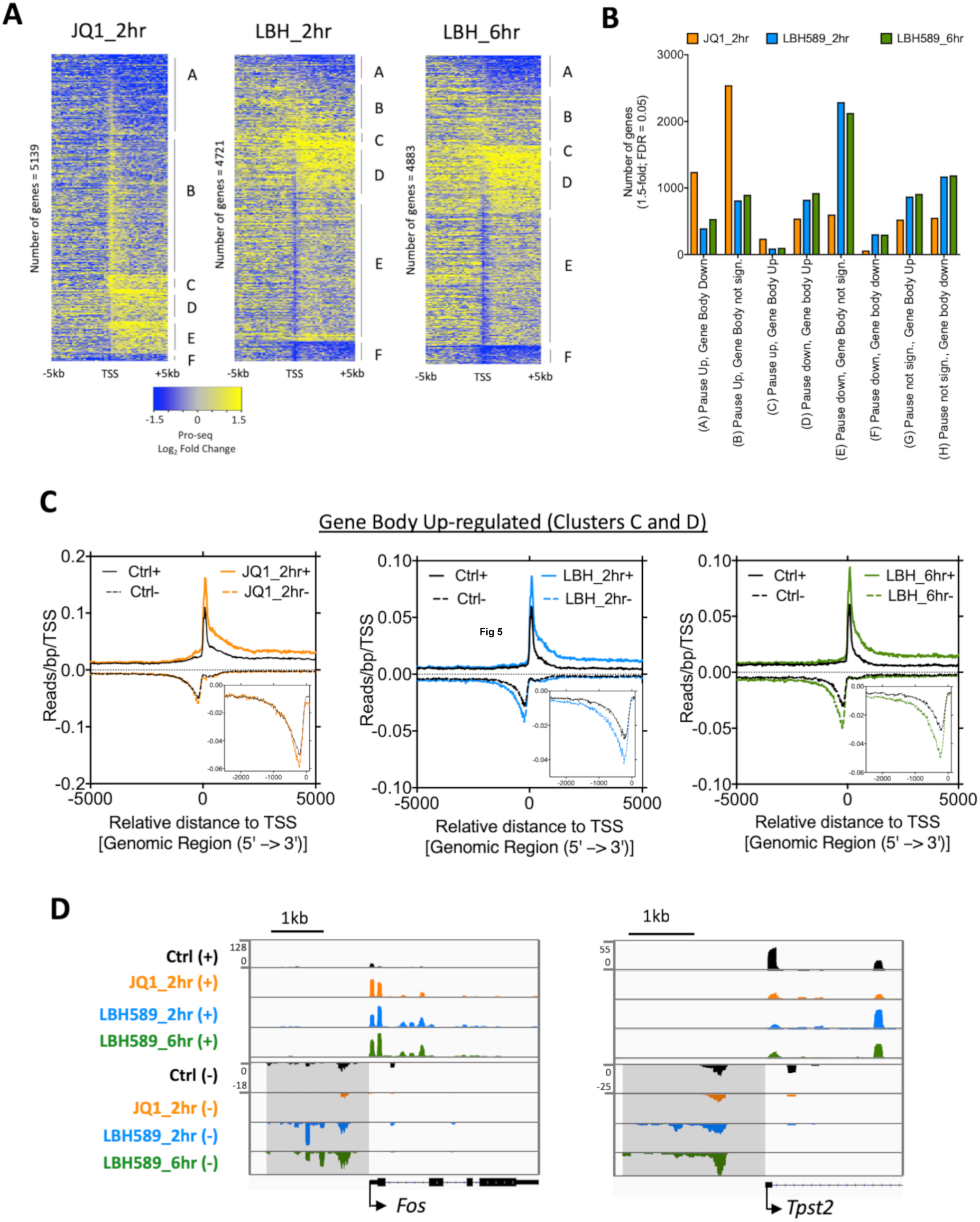
HDAC inhibition caused increased expression of promoter upstream transcripts (PROMPTs) and a loss of promotor proximal polymerase. (A) Heatmaps displaying a log_2_-fold change of PRO-seq read counts in 200-bp bins (TSS± 5 kb) at Refseq genes displaying increased or decreased pause index changes following JQ1 (2hr) or LBH589 (2hr and 6hr) treatment. Genes were ranked based on log_2_-transformed fold-change of RNA polymerase in the promoter-proximal region. (B) A number of gene clusters (A-F) with similar transcriptional changes are shown. For each treatment genes were clustered based on (± 1.5 fold; FDR≤0.05) change in gene body or paused index changes. (C) Metagene plots (TSS ± 5 kb) of RNA polymerase densities at genes showing increased gene body levels following the treatment with JQ1 or LBH589. Inserts show increased antisense transcript (PROMPTs) relative to TSS in greater detail. (D) PRO-seq IGV bedGraph screenshots of *Fos* and *Tpst2* TSS in control with JQ1 or LBH589 for the indicated times. Shaded areas indicate increased antisense transcription in LBH589-treated samples.

### HDAC activity is required for RNA polymerase recruitment to pluripotent genes

Since both LBH589 and JQ1 attenuate *Nanog*, *Pou5f1* and *Sox2* expression (Figure 4A), we used our PRO-seq data to assess their roles in transcriptional pausing and elongation among the wider network of factors required for pluripotency. Transcription was reduced in 34 of 59 (2 hr) and 45 of 59 (6 hr) pluripotent network genes in LBH589-treated cells and 30 of 59 in cells with JQ1 (Figure 6A), demonstrating a direct requirement for both deacetylase activity and BET family members. The reduction in transcription following JQ1 treatment is accompanied by a significant increase in Paused Index (PI, the ratio of promoter-proximal to gene body transcript levels (Kwak et al. 2013)(Figure 6B), indicating that reduced expression is due to an inability to resolve paused RNA pol II. In contrast, HDAC inhibition produced no consistent change in PI among pluripotent factors (Figure 6B, compare JQ1 with LBH589 at 2 and 6 hours). Metagene plots of polymerase density at pluripotent genes (Figure 6C) showed increased RNA Pol II pausing with JQ1, while LBH589 treatment caused a reduction in both promoter-proximal and gene body transcription. A reduction in both paused and elongating polymerase can be observed at individual genes such as *Nanog*, Sox2, *Pou5f1*, *Klf2* and *Sall4* (Figure 6D). An overall reduction in transcription suggests that HDAC activity may be required for optimal RNA Pol II recruitment. To test this further, we performed ChIP-qPCR for RNA Pol II at the *Nanog*, *Klf4* and *Pou5f1* promoters in the presence or absence of LBH589 (Figure 6E). We observed reduced RNA pol II binding at all three genes when treated with LBH589 compared to controls, but intriguingly, there was no change with JQ1. Taken together, these findings demonstrate that HDAC inhibition reduced the level of both paused and elongating polymerase at pluripotent genes, consistent with a role in RNA Pol II recruitment that appears to be mechanistically independent of BRD4.

**Figure 6:**
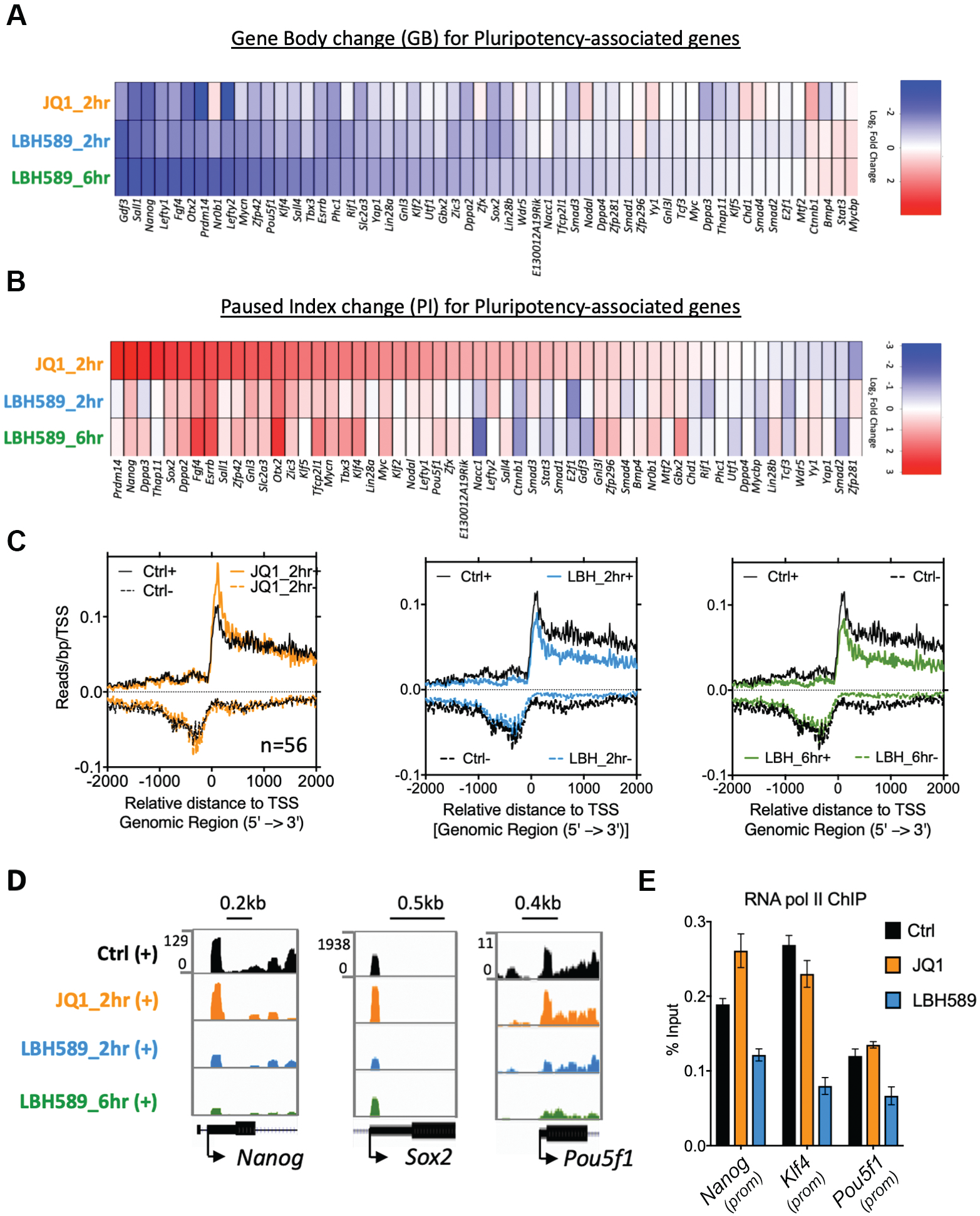
HDACs regulate expression of the pluripotent gene network via recruitment of RNA polymerase II. (A) PRO-seq analysis displaying the effects of JQ1 or LBH589 treatment on the level of gene body transcripts for pluripotent gene network members. Heatmap shows log_2_ fold-change relative to untreated controls. (B) Heatmap shows the effects of JQ1 and LBH589 treatment on pausing index (the ratio of promoter-proximal to gene body transcript levels) changes relative to untreated controls. (C) Metagene plots (TSS ± 4 kb) of pluripotency genes demonstrating decreased initiation and elongation in LBH589 treated samples. (D) RNA polymerase density at *Nanog, Sox2* and *Pou5f1, Klf2 and Sall4* TSS in either control, JQ1 or LBH589 treated ESCs.

### HDACs positively regulate a subset of enhancers associated with the pluripotent network

Analysis of PRO-seq data using nascent RNA sequencing analysis (NRSA) (Wang et al. 2018) identified transcripts in intergenic regions corresponding to enhancer RNAs (eRNAs). Since eRNA levels are a more sensitive indicator of enhancer activity than the histone modifications (Kim et al. 2010), we can directly infer the effects of LBH589 and JQ1 on enhancer activity from their transcription. Consistent with changes in mRNA levels (Figure 3A), we observed a similar number of down-(199) and up-regulated (149) eRNAs following a 2-hour LBH589 treatment, with an increasing number (627 down and 539 up) at 6 hours (Figure 7A). In contrast, the addition of JQ1 predominantly down-regulated eRNAs (1196 down vs 33 up) and overall affected a far greater number of eRNAs than LBH589. These data indicate that BET family members play an important role in the expression of eRNAs and that this is independent of HDAC activity.

**Figure 7:**
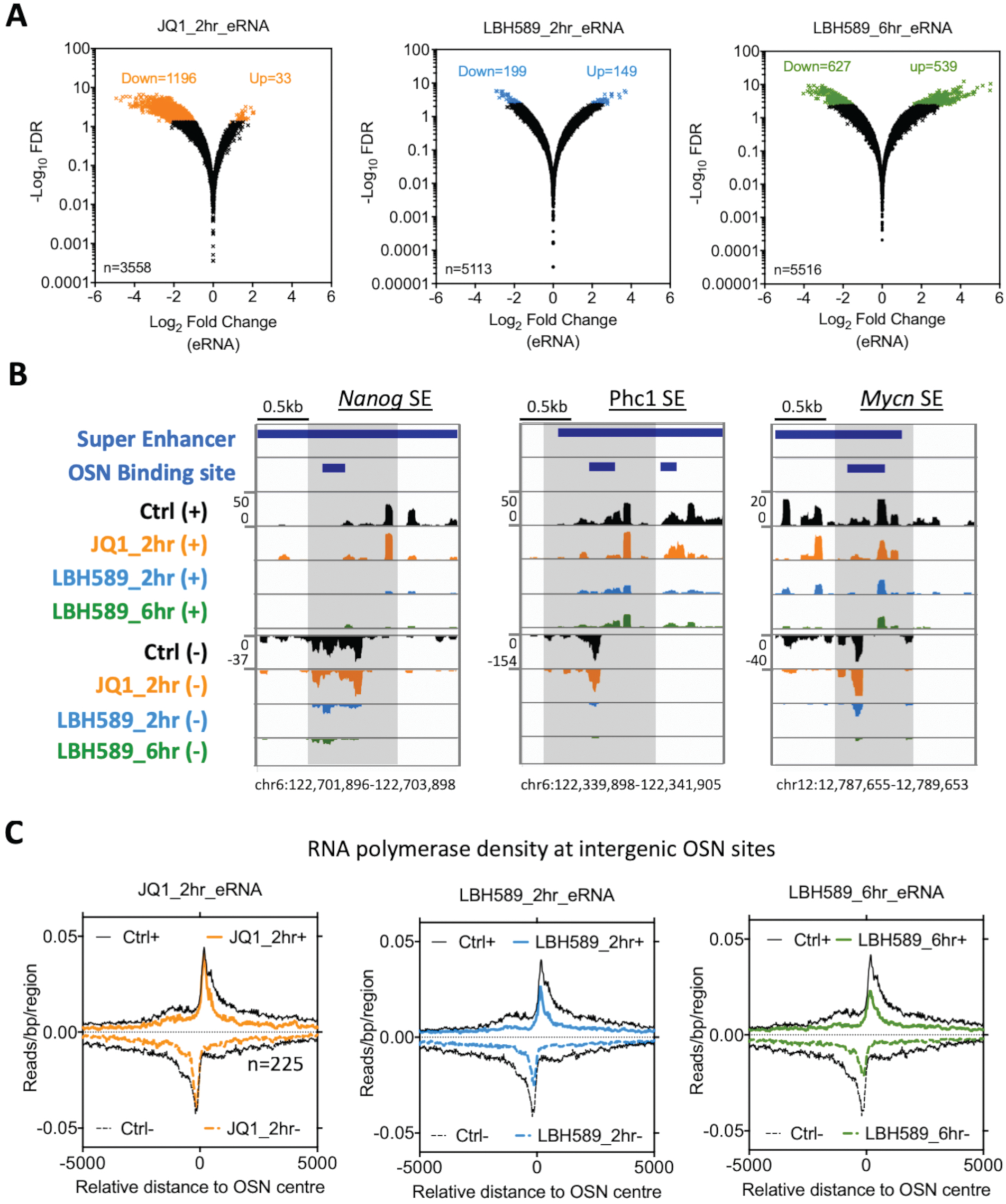
HDACs positively regulate a subset of eRNAs associated with super-enhancers. (A) Volcano plot showing changes in eRNA transcription following treatment with JQ1 or LBH589 for the indicated time points. Statistically enriched (± 1.5 fold; FDR≤0.05) eRNA transcripts relative are indicated by colour. (B) IGV genomic tracks of PRO-seq RNA polymerase densities at intergenic OCT4/SOX2/NANOG (OSN) binding sites associated with the indicated super-enhancer regions. Shaded areas indicate regions of decreased transcription following LBH589 treatment. (C) Metagene plots of RNA polymerase density reads at intergenic OSN binding sites following the indicated treatments.

Both BRD4 (Di Micco et al. 2014) and HDACs (Sanchez et al. 2018) are known to alter transcriptional dynamics at SEs. Therefore, we examined the transcription of eRNAs that overlap with SEs in ESCs treated with either LBH589 or JQ1. Intriguingly, a greater proportion of the eRNAs down-regulated by LBH589 overlapped with SEs (39 of 199 – 20%) than those with JQ1 (57 of 1199 - 5%)(Figure 7A). Examination of individual SEs (*Nanog*, *Mycn* and *Phc1*) shows a distinct reduction in transcription upon HDAC inhibition that appeared unaffected by JQ1 treatment (Figure 7B, compare orange vs blue/green peaks in shaded regions). Since SEs are defined in part by increased recruitment of lineage-specific transcription factors, we hypothesized that HDAC activity may be required for eRNAs associated with the core pluripotent transcription factors, *Oct4 (Pouf51), Sox2* and *Nanog* (OSN). We performed an unbiased analysis of eRNA transcription at all OSN binding sites. Upon HDAC inhibition, we observed a reduction in RNA pol II density at both the TSS and the body of the transcript indicating a reduction in recruitment, while JQ1 predominantly affected elongation (Figure 7C). These data suggest that within 2 hrs of LBH589 treatment, enhancers associated with OSN function have reduced activity, contributing to the decrease in expression of the entire pluripotent network (Figure 6A).

## Discussion

HDAC enzymes and their associated complexes (e.g. Sin3A) are generally viewed as negative regulators of gene expression. The removal of acetyl groups from histone tails leads to a tightening of chromatin that prevents binding of transcription factors. Moreover, transcriptionally silent regions of the genome generally contain hypo-acetylated histones (Barnes et al. 2019) and references therein). Despite this, there are several lines of evidence that indicate that HDACs play a role in active transcription. ChIP experiments for class I HDACs indicate that they are recruited to both enhancers and promoters of active genes (Wang et al. 2009; Kidder and Palmer 2012; Whyte et al. 2012). Acetylation/deacetylation cycling may be required to reset promoters between rounds of initiation (Metivier et al. 2003). The latter results have only been demonstrated in a limited number of genes, but the concept is supported by the dynamic turnover of histone acetylation. Acetylation sites within the H2B tail in particular are subject to rapid deacetylation with a half-life of approximately 10 min (Weinert et al. 2018). Although these data all support a role in transcription, the readout of histone acetylation remains highly dependent on genomic context, since HDAC inhibition results in both up-and down-regulated genes (Dovey et al. 2010a).

In ESCs, we have shown that HDAC1/2 (Figure 1A), but not HDAC3 (Supplementary Figure 1C), are required for the expression of the pluripotent network. This activity is dependent on gene dosage since a hypomorphic *Hdac1-KO; Hdac2-Het* cell line has an intermediate level of *Pou5f1* and *Nanog* expression compared to single (Dovey et al. 2010b), or DKO ESCs (Figure 1A and (Jamaladdin et al. 2014)). HDAC1/2 are incorporated into six biochemically and functionally distinct multi-protein complexes: SIN3, NuRD, CoREST, MiDAC, MIER and RERE (Kelly and Cowley 2013; Millard et al. 2017). ESCs in which the NuRD (Kaji et al. 2006; Burgold et al. 2019), CoREST (Foster et al. 2010) and MiDAC (Turnbull et al. 2020) complexes are perturbed still retain pluripotency. In contrast, knock-down of SIN3A (Saunders et al. 2006; Fazzio et al. 2008; Baltus et al. 2009; Adams et al. 2018; Zhu et al. 2018), or SIN3A complex components (Streubel et al. 2017) reduced the expression of *Nanog* and other pluripotent factors, suggesting that the Sin3 complex may be the critical HDAC1/2 complex component.

Although *Hdac1/2* double knock-out ESCs show a robust decrease in *Nanog* transcription (Figure 1A), the stable nature of class I HDACs (half-life of 24 hours) obliged us to use HDACi to examine the direct effects of deacetylation on transcription. Analysis of nascent transcripts in ESCs showed a significant reduction in *Nanog* and *Pou5f1* transcription within 2 hours of LBH589 treatment (Figure 3E). The similarity between LBH589 treatment (a general inhibitor of Zn^2+^-dependent HDACs) and *Hdac1/2* DKO cells, suggests that this acute decrease in pluripotent factors is a result of HDAC1/2 inhibition. A critical role of HDAC1/2 in ESCs is to regulate histone acetylation levels (e.g. H3K18ac) across the genome (Kelly et al. 2018), thereby indirectly affecting the recruitment of acetyl-lysine readers, such as BRD4. Increased histone acetylation across the genome could potentially sequester a limiting pool of BRD4 from critical promoters, resulting in a reduced transcription (Greer et al. 2015). In ESCs, we find that over-expression of BRD4 increased the level of *Nanog* expression indicating that it is limiting (Figure 4C). However, overexpression of BRD4 was insufficient to rescue the decrease in Nanog transcription upon HDAC inhibition, arguing in favour of the collaborative but independent roles of BRD4 and HDACs.

To examine the direct roles of HDACs and BET family members (BRD2-4), we performed PRO-seq in ESCs treated with both LBH589 and the bromodomain inhibitor, JQ1 (Figure 5). PRO-seq allows high-resolution mapping of active RNA polymerases across the genome at both coding and non-coding regions and provides information on both the abundance and directionality of transcripts. Analysis of the transcriptome in LBH589-treated ESCs revealed a reduction in both promoter-proximal RNA pol II *and* elongating polymerase within the gene body (Figure 4A), indicating an overall reduction in recruitment. Indeed, ChIP-qPCR confirmed a reduction in RNA Pol II recruitment at the *Nanog*, *Pou5f1* and *Klf4* promoters following HDAC inhibition (Figure 6E). This contrasts with the addition of JQ1, which produced a significant increase in paused polymerase (Figure 5A), but no alteration in Pol II recruitment (Figure 6E).

RNA Pol II recruitment to promoters is the principal function of enhancers. Since enhancer activity is associated with the production of eRNAs, whose levels are correlated with the activity of neighbouring genes (Andersson et al. 2014; Core et al. 2014), we used our PRO-seq data to examine the contribution of HDACs and BET family members to enhancer function in ESCs (Figure 7A). JQ1 treatment reduced the transcription of 1196 eRNAs, compared to only 199 with LBH589. However, the LBH589-sensitive eRNAs contained a far greater proportion of eRNAs associated with super-enhancer (SE) activity (Figure 7B). The directionality of transcripts identified by PRO-seq also allows the identification of sub-enhancers contained with SEs. Examination of individual SEs, such as the *Nanog*, *Phc1* and *Mycn*, demonstrates an overall reduction in transcripts and RNA Pol II recruitment at each of the sub-enhancers within these regions (Figure 7B).

While our study has identified a role for HDACs in the activity of SEs in ESC, there is accumulating evidence that this might represent a broader mechanism in other cell types. Treatment of the human colon tumour cell line HCT116 with the natural HDACi, largazole, resulted in a reduction in RNA Pol II recruitment to the *Myc* SE (Sanchez et al. 2018). Intriguingly, reduced polymerase recruitment was independent of H3K27ac levels, although it remains possible that alternative sites of histone acetylation might be involved (Sanchez et al. 2018). Gryder et al. used constitutive and SE-driven reporters to identify classes of histone-modifying enzymes that preferentially affect SE-driven transcription in alveolar rhabdomyosarcoma cells. Remarkably, they found that SEs were more sensitive to HDAC inhibitors than a range of bromodomain inhibitors, including JQ1 (Gryder et al. 2019b). Subsequently, they showed that HDAC inhibition triggers hyperacetylation-induced erosion of SE boundaries which alters 3D chromatin dynamics and reduces RNA Pol II occupancy at specific gene loci (Gryder et al. 2019a). In ESCs, we observed an analogous increase in H3K27ac levels at SEs and reduced RNA Pol II occupancy upon HDAC inhibition, providing evidence of a broader mechanistic requirement for HDAC activity in SE function.

Transcription factors that define a cell-specific transcriptional programme (such as Oct4 and Nanog) have increased clusters of binding sites at cell-type-specific SEs, which in turn, regulate their own expression (Whyte et al. 2013). This self-reinforcing auto-regulatory network creates a system whereby loss of activity (e.g. treatment with HDACi) leads to a ‘self-reinforcing’ collapse (Gryder et al. 2019b). This hypothesis would fit with our data in ESCs, which showed a reduction in eRNAs associated with OSN binding sites following LBH589 treatment (Figure 7C). And a collective reduction in OSN activity would therefore lead to a general loss of expression for the entire pluripotent network (Figure 6A).

## Data Availability

All microarray and sequencing data have been deposited in the GEO database under the SuperSeries GSE136621. Cell lines and plasmids used in the study are available upon request.

## Acknowledgements

We thank Prof Ian Eperon, Dr Grace Adams, India-May Baker and Donovan Lim for critical reading of the manuscript. We are grateful to Dr Nicolas Sylvius and the NUCLEUS Genomics facility for help with performing the microarray data collection; data analysis was assisted by Dr Matthew Blades. Plasmid constructs were generated by the University of Leicester PROTEX facility.

