## Supplementary Fig 1, Tables 1 and 2 for "Histone Deacetylases (HDACs) maintain expression of the pluripotent gene network via recruitment of RNA polymerase II to coding and non-coding loci"

**Supplemental Figure 1**

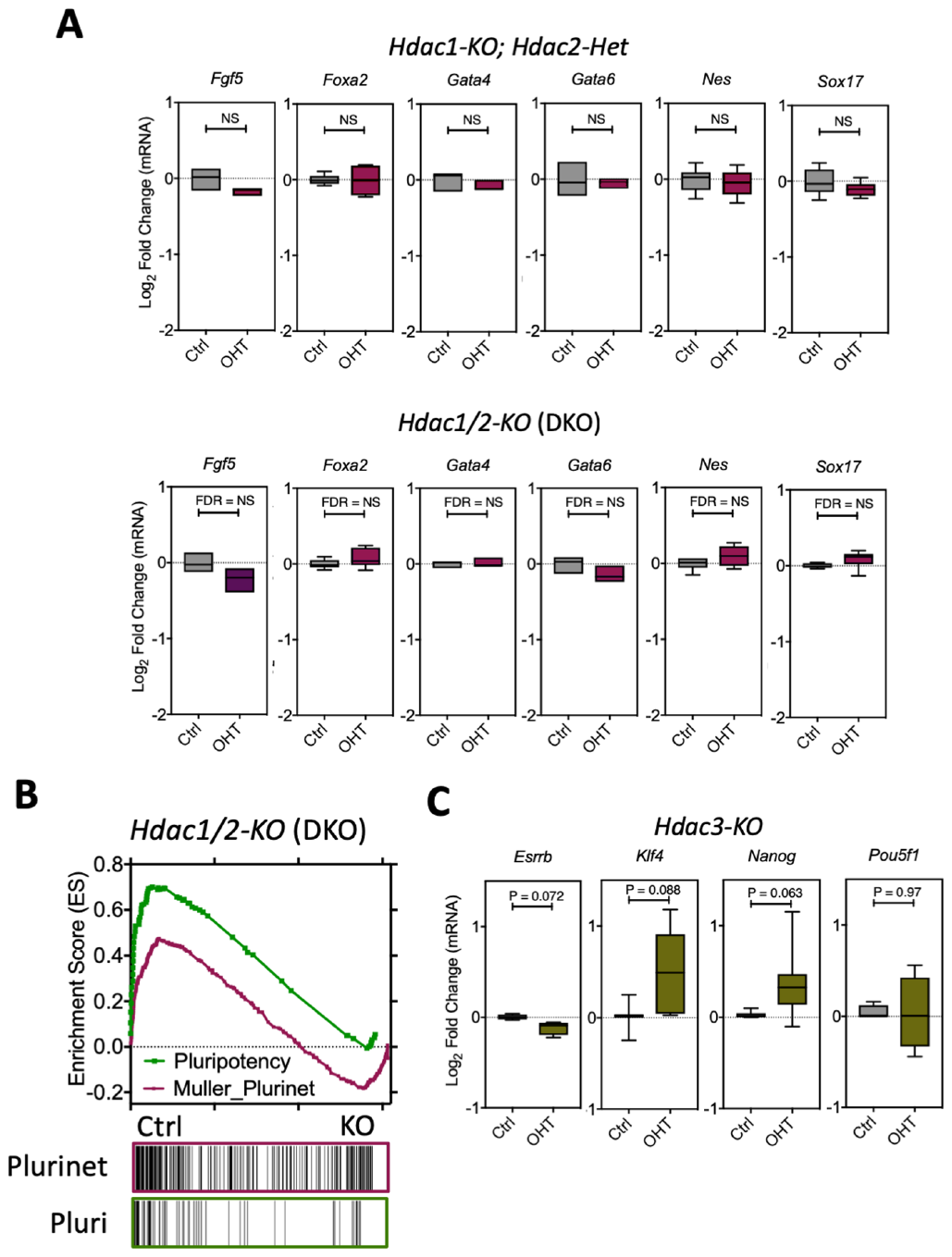

**Supplemental Fig 1 – Loss of HDAC1/2 or HDAC3 does not lead to differentiation of embryonic stem cells (ESC).** (A) Box plots generated form microarray data from compound (*Hdac1-KO; Hdac2-Het*) or double knockout (DKO) ESCs, treated with 4-hyrdoxytamoxifen (OHT) or and untreated control (Ctrl). Microarray data from n = 3 biological replicates were used to generate floating box plots (min-max) with line at mean. Statistical differences were calculated using Benjamini & Hochberg false discovery rate (FDR). (B) Gene set enrichment analysis (GSEA) plot of microarray data showing Ctrl samples are enriched for the Muller Plurinet (ES=0.47; P<0.0001) and pluripotency (ES=0.70; P<0.0001) gene sets compared to DKO cells. (C) Box plots generated form microarray data from *Hdac3-KO* ESCs, treated with 4-hyrdoxytamoxifen (OHT) or and untreated control (Ctrl). Microarray data from n = 3 biological replicates were used to generate floating box plots (min-max) with line at mean. Statistical differences were calculated using Benjamini & Hochberg false discovery rate (FDR).

**Supplementary Table 1**

| **Antibody** | **Source** | **Clonality** | **Company** | **Application** | **Concentration** |
| --- | --- | --- | --- | --- | --- |
| HDAC1 | Rabbit | Monoclonal | Abcam (ab109411) | Western Blotting | 1:2500 |
| HDAC2 | Mouse | Monoclonal | Millipore (05-814) | Western Blotting | 1:2000 |
| NANOG | Rabbit | Polyclonal | Bethyl (A300-397A) | Western Blotting | 1:2500 |
| POU5F1/OCT4 | Mouse | Monoclonal | Santa Cruz (SC5279) | Western Blotting | 1:500 |
| 𝛼-Tubulin | Mouse | Monoclonal | Millipore/Sigma (T5168) | Western Blotting | 1:15000 |
| BRD4 | Mouse | Polyclonal | Bethyl (A301-985A) | ChIP-qPCR | 10 μL /ChIP |
| H3K27ac | Rabbit | Polyclonal | Active Motif (39135) | ChIP-qPCR | 8 μg /ChIP |
| H3K56ac | Rabbit | Polyclonal | Active Motif (39281) | ChIP-qPCR | 10 μL/ChIP |
| RNA polymerase II | Mouse | Monoclonal | Active Motif (39097) | ChIP-qPCR | 10 μL /ChIP |

**Supplementary Table 2**

| **Accession Number** | **Target** | **Primer Name** | **Sequence** | **Annealing temp** | **Application** |
| --- | --- | --- | --- | --- | --- |
| NM_011934 | *Esrrb* | Esrrb_R | CATGAAATGCCTCAAAGTGGG | 58°C | RT-qPCR |
|  |  | Esrrb_F | AAATCGGCAGGTTCAGGTAG |  |  |
| NM_010637 | *Klf4* | Klf4_F | TGTGTCGGAGGAAGAGGAAGC | 59°C | RT-qPCR |
|  |  | Klf4_R | ACGACTCACCAAGCACCATCA |  |  |
| NM_028016 | *Nanog* | Nanog_F | AGGGTCTGCTACTGAGATGCTCTG | 60°C | RT-qPCR |
|  |  | Nanog_R | CAACCACTGGTTTTTCTGCCACCG |  |  |
| NM_013633 | *Pou5f1* | Pou5f1_F | AGTATGAGGCTACAGGGACA | 60°C | RT-qPCR |
|  |  | Pou5f1_R | CAAAGCTCCAGGTTCTCTTG |  |  |
| NM_023755 | *Tfcp2l1* | Tfcp2l1_F | AGGTGCTGACCTCCTGAAGA | 58°C | RT-qPCR |
|  |  | Tfcp2l1_R | GTTTTGCTCCAGCTCCTGAC |  |  |
| NM_009382 | *Thy1* | Thy1_R | TGCTCTCAGTCTTGCAGGTG | 57°C | RT-qPCR |
|  |  | Thy1_F | TGGATGGAGTTATCCTTGGTGTT |  |  |
| NM_139218 | *Dppa3* | Dppa3_F | TCGACCCAATGAAGGACCCTGAAA | 60°C | RT-qPCR |
|  |  | Dppa3_R | TTGGGAAAGGCGCTTTGAACTTCC |  |  |
| NM_008108 | *Gdf3* | Gdf3_F | TAAGGATTGGAGCAGCAACCGACT | 60°C | RT-qPCR |
|  |  | Gdf3_R | ATGACGGTGGCAGAAGTTCCTACA |  |  |
| NM_011443 | *Sox2* | Sox2_F | TAGAGCTAGACTCCGGGCGATGA | 60°C | RT-qPCR |
|  |  | Sox2_R | TTGCCTTAAACAAGACCACGAAA |  |  |
| NM_007393 | *β-ACTIN* | b_Actin_F | GGCTCCTAGCACCATGAAGA | 60°C | RT-qPCR |
|  |  | b_Actin_R | AGCTCAGTAACAGTCCGCCT |  |  |
| NC_000072 | *Nanog*  *enhancer (5kb)* | Nanog_enhancer (5kb)_F | GTTTTGACTGCTAACCACCCAGAG | 58°C | ChIP_qPCR |
|  |  | Nanog_enhancer (5kb)_R | GGCAGGCTTGCTACATTCCTTATC |  |  |
| NC_000072 | *Nanog*  *promoter* | Nanog_promoter_F | ACAGCTTCTTTTGCATTACAATGTCC | 58°C | ChIP_qPCR |
|  |  | Nanog_promoter_R | TATTCTCCCAGGCACCCAGGC |  |  |
| NC_000083 | *Pou5f1*  *enhancer (-2kb)* | Pou5f1_enhancer (-2kb)_F | AGGTGCATGATAGCTCTGCC | 58°C | ChIP_qPCR |
|  |  | Pou5f1_enhancer (-2kb)_R | AAGCCGCCAAGTTCACAAAG |  |  |
| NC_000072 | *Gata2*  *promoter* | Gata2_promoter_F | GGTCATTCCCGAAGTCCAGC | 58°C | ChIP_qPCR |
|  |  | Gata2_promoter_R | GACTCCTGCACAGACGTGAA |  |  |
| NC_000071 | *Spink2*  *promoter* | Spink2_promoter_F | CCCCAAACACATCCTTTCGAC | 58°C | ChIP_qPCR |
|  |  | Spink2_promoter_R | TATGGAGTCTGTCGGGGGAA |  |  |
| NC_000067 | *Tfcp2l1*  *enhancer (-2kb)* | Tfcp2l1_enhancer (-2kb)_F | AATCAAAAGGCTGCCCTTGC | 58°C | ChIP_qPCR |
|  |  | Tfcp2l1_enhancer (-2kb)_R | TCCCGGCCCTAAGAACCTAA |  |  |
| NC_000072 | *Nanog*  *TSS* | Nanog_TSS_F | GTGCAGCCGTGGTTAAAAG | 58°C | ChIP_qPCR |
|  |  | Nanog_TSS_R | GGTCCACCATGGACATTGT |  |  |
| NC_000070 | *Klf4 TSS* | Klf4_TSS_F | TAACTTCTCGCTCGCTTGCT | 58°C | ChIP_qPCR |
|  |  | Klf4_TSS_R | CGTGCGCGGAGTTTGTTTAT |  |  |
| NC_000083 | *Pou5f1*  *TSS* | Pou5f1_TSS_F | GATCCTCGAACCTGGCTAAG | 58°C | ChIP_qPCR |
|  |  | Pou5f1_TSS_R | CAGGCTGCAAAGTCTCCAC |  |  |
